# Interpretable Machine Learning Decodes Soil Microbiome’s Response to Drought Stress

**DOI:** 10.1101/2023.11.30.569182

**Authors:** Michelle Hagen, Rupashree Dass, Cathy Westhues, Jochen Blom, Sebastian J Schultheiss, Sascha Patz

**Affiliations:** Computomics GmbH, Eisenbahnstraße 1, Tübingen, 72072, Baden-Württemberg, Germany; Bioinformatics & Systems Biology, Justus Liebig University Gießen, Heinrich-Buff-Ring 58, Gießen, 35390, Hesse, Germany

**Keywords:** Metagenomics, Machine Learning, SHAP values, Differential Abundance Analysis, Soil microbiome, Drought stress

## Abstract

**Background:** Extreme weather events induced by climate change, particularly droughts, have detrimental consequences for crop yields and food security. Concurrently, these conditions provoke substantial changes in the soil metagenome and affect plant health. Early recognition of soil affected by drought enables farmers to implement appropriate agricultural management practices. In this context, interpretable Machine Learning holds immense potential for drought stress classification in the soil metagenome based on marker taxa.

**Results:** This study demonstrates that the metagenomic approach of Differential Abundance Analysis methods and Machine Learning-based Shapley Additive Explanation values provide similar information. They exhibit their potential as complementary approaches for identifying marker taxa and investigating their enrichment or depletion under drought stress in grass lineages. Additionally, the Random Forest Classifier trained on a diverse range of relative abundance data from the soil metagenome of various plant species achieves a high accuracy of 92.3 % at the genus rank for drought stress prediction. It demonstrates its generalization capacity for the lineages tested.

**Conclusions:** In the detection of drought stress in the soil metagenome, this study emphasizes the potential of an optimized and generalized location-based ML classifier. By identifying marker taxa, this approach holds promising implications for microbe-assisted plant breeding programs and contributes to the development of sustainable agriculture practices. These findings are crucial for preserving global food security in the face of climate change.

## 1 Background

Global food security is significantly threatened by climate change, especially in regions with limited access to food resources [1–3]. Anticipated occurrences of extreme weather events, such as droughts, are likely to increase in frequency and intensity, causing significant crop damage and threatening food availability [4, 5]. These repercussions are attributed to shortened growing seasons and substantial reductions in crop yields due to various biotic and abiotic stresses, prominently drought stress [6–8].

External perturbations, like drought stress, significantly impact the dynamics of the soil microbial community, leading to compositional shifts that are facilitated by the recruitment of beneficial microbes from the surrounding soil to the roots [9, 10]. This interaction between plants and soil microorganisms is a vital aspect of ecosystem health and stability [11–14]. Hence, this results in the opportunity to identify and interpret specific metagenomic patterns, as they have the potential to provide valuable insights into the state of both soil and plant health.

The use of Machine Learning (ML) algorithms enables the analysis of complex microbiome data by fully capturing the depth of data and identifying patterns that can discriminate between different states or conditions [15]. This can help to identify specific marker taxa that are key to understanding the intricate relationships between environmental stressors, soil health, and plant viability. Thereby, they aid in the development of more effective soil management strategies, including irrigation practices and the selection of drought-tolerant crops. Marker taxa hold significant potential for application in Synthetic Communities (SynComs), providing an innovative approach for early intervention under challenging environmental conditions, like droughts, to enhance plant resilience and growth [10, 16].

Still, the path from ML predictions to actionable insights can be challenging. ML models often resemble black boxes, with their internal decision-making obscured from users. The interpretation of the reasons for certain predictions is essential, especially for complex biological data [17–19]. This is where interpretable ML methods such as SHapley Additive ExPlanation (SHAP) values are applied [20].

The concept behind SHAP values is to distribute the credit for the model’s prediction among the feature inputs based on their individual contribution, using game theory. Notably, SHAP values are model-specific yet globally constant, comprehensively taking into account interactions between features [21].

While SHAP values have been applied in clinical studies [22, 23], their use in metagenomic data is limited. For the identification of significant taxa between comparison groups, Differential Abundance Analysis (DAA) is the commonly used method in metagenomic analyses [24]. Typically used DAA tools are DESeq2 [25], ALDEx2 [26], edgeR [27], ANCOM-BC2 [28], and the non-parametric Wilcoxon rank-sum test. DAA tools and SHAP values offer distinct approaches for detecting marker taxa.

DAA tools rely on statistical tests and assume specific data distributions [29], while SHAP values are model-agnostic and applicable to any ML model [21]. Both methods aim to identify crucial features or taxa, providing insights into underlying biological mechanisms, but serve different purposes. DAA methods focus on identifying differentially abundant taxa, whereas interpretable ML using SHAP values offers importance measures based on model performance. The main objective of SHAP values is to interpret complex ML models by quantifying the contribution of each feature and explaining predictions. Their application in this study demonstrates the potential to identify key taxa in soil microbiomes as well as their role in the microbial response to drought stress.

The selection of an appropriate soil dataset was essential for this study. ML analyses thrive on datasets with many samples and informative metadata [30]. A dataset from the work of *Naylor et al*. [31] was selected because it met these criteria exceptionally well. This dataset includes 623 samples from three soil isolation sources and investigates the effect of drought stress on 19 different crop species, including C3 and C4 plants.

Despite the remarkable achievements in applying ML to human microbiome research [32–36], its application in the context of soil metagenomics is not yet as advanced. However, the agricultural industry is increasingly recognizing the potential of ML to improve soil health and promote sustainable farming practices [37, 38]. This includes predicting plant phenotypes based on the plant and surrounding soil microbiome to detect taxa associated with plant diseases and environmental stresses [39].

Prior research has highlighted the potential of ML in agriculture, with studies identifying marker taxa for crop productivity [40] and beneficial root microbes [41]. Interestingly, the potential of ML for drought stress identification in soil microbiomes remains largely unstudied, representing a promising area for investigation.

This research aims to determine the efficacy of ML in predicting drought stress within microbial data of drought-stressed soils. The study comprises three key objectives: a) investigating the predictive capability of ML for drought stress, b) comparing the performance of interpretable ML with conventional 16S rRNA-based metagenomic analyses, and c) assessing the generalization capabilities of the trained classifier. By identifying marker taxa and deciphering microbial patterns associated with drought stress, this research addresses sustainable agriculture, improved crop productivity, and increased food security.

## 2 Methods

### 2.1 Datasets

A dataset originally curated by *Naylor et al*. [31] for their study on the impact of drought stress on the grass root microbiome was analyzed. This dataset, referred to as the ‘Grass-Drought’ dataset, comprises 623 samples from three isolation sources, including ‘Soil,’ ‘Root,’ and ‘Rhizosphere’, as well as two watering regimes, including ‘Drought’ and ‘Control’. Samples from 18 distinct grass species within the Poaceae clade are included in the dataset. Tomato was used as an outgroup. The experimental site was located in Albany, California, characterized by silty loam soil with a pH of 5.2. Both watering regimes, ‘Drought’ and ‘Control’, were balanced, with 320 samples in the ‘Control’ group, receiving regular watering, and 303 samples in the ‘Drought’ group, experiencing conditions without water supply. All samples were sequenced using 16S rRNA amplicon sequencing of the V3-V4 region and are available under the BioProjectID PRJNA369551.

To evaluate the ML model’s generalizability, its performance was assessed on a separate test dataset from *Xu et al*. [42] studying pre- and post-flowering drought stress effects on the *Sorghum bicolor* root microbiome (BioProjectID PRJNA435634), therefore referred to as the ‘Sorghum-Drought’ dataset. The sampling site was located in Kearney, California. To ensure the comparability of drought conditions between the Sorghum-Drought dataset and the original Grass-Drought dataset, two subsets were created: The ‘Progressive Drought’ subset comprised samples from the ‘Control’ group, along with specific time points (weeks 2 to 7 and weeks 10 to 17) from the ‘Pre-Flowering Drought’ and ‘Post-Flowering Drought’ groups, respectively. This subset comprised 278 ‘Control’ and 210 ‘Drought’ samples. The ‘Late Drought’ subset included samples from weeks 6, 7, 16, and 17 of the ‘Control’ group, weeks 6 and 7 of the ‘Pre-Flowering Drought’ group, and weeks 16 and 17 of the ‘Post-Flowering Drought’ group, totaling 72 ‘Control’ and 69 ‘Drought’ samples. A detailed representation of the subsetting scheme can be found in the Additional file 1 in **Fig. S1**.

### 2.2 Data Processing

The DADA2 workflow [43] for Illumina sequenced paired-end fastq files was employed for sequence data processing, implemented in R version 4.2.3. Taxonomy was assigned using the SILVA database [44] and the Ribosomal Database Project (RDP) classifier [45] from phylum to genus rank. To enhance data quality, prevalence filtering was conducted, retaining Amplicon Sequence Variants (ASVs) present in at least 95 % of all samples, reducing the total number of ASVs from 25,415 to 3,276. Samples with low read counts were excluded, yielding a dataset of 560 samples. Rarefaction was performed, normalizing sequencing depth to the dataset’s 10 % decile of 17,291 reads. Feature tables for ML for each taxonomic rank were constructed with relative abundance values per taxon across all samples and a ‘Control’ or ‘Drought’ target variable.

### 2.3 16S rRNA-based Metagenomic Analysis

A diversity analysis was conducted between the two watering regimes ‘Control’ and ‘Drought’. Alpha diversity was assessed using the Shannon index with the estimate richness function from the phyloseq package (version 1.44.0) [46]. Beta diversity was explored via Principal Coordinate Analysis (PCoA) based on Bray-Curtis dissimilarities with the ordinate and plot ordination functions from phyloseq.

To identify taxonomic differences between the ‘Control’ and ‘Drought’ groups, a DAA was employed with several tools using the microbiomeMarker R package (version 1.4.0) [47], including DESeq2, ALDEx2, and edgeR, as well as ANCOM-BC2 (version 2.0.2) [28], and the non-parametric Wilcoxon rank-sum test on the ASV level. The UpSetR package (version 1.4.0) [48] was used to compare tool outcomes, and the three most suitable methods were applied to all taxonomic ranks to compare enrichment groups and Benjamini-Hochberg (BH) adjusted p-values [49].

### 2.4 Machine Learning

A Random Forest Classifier (RFC) was applied using the ScikitLearn Python package (version 1.1.3) [50] to all ranks to predict the samples’ watering treatment using relative taxon abundances. Hyperparameter optimization was carried out through five-fold nested Cross-Validation (CV), splitting the dataset into five equally sized parts. During each fold, four parts of the dataset were used for training, while the remaining part acted as a dataset for testing the best model of each fold. The mean model performance was evaluated in terms of accuracy, F1 score, precision, recall, and Area Under the Curve (AUC) between all folds.

In order to interpret the RFC predictions, SHAP values were utilized using the SHAP Python package (version 0.41.0) [51] with the shap.TreeExplainer function [20]. During each fold of the nested cross-validation, feature contributions related to detecting drought stress from SHAP values were extracted. A consensus was sought across four or five of the folds, requiring alignment in the majority, to consider the enrichment information suitable for subsequent analysis. The feature contributions towards drought stress from the SHAP values were compared with taxon enrichment patterns from differential abundance testing. This was followed by a comparison of significant taxa identified by DAA methods and important taxa identified by ML.

The model performance and generalizability were tested on two independent subsets of the Sorghum-Drought dataset of *Xu et al*. [42] that has been described in detail in the ‘Datasets’ section. This test dataset was processed similarly to the Grass-Drought dataset. To create feature tables, the tables were pruned to only include taxa present in the Grass-Drought dataset. Taxa that were not present in the Sorghum-Drought dataset but in the Grass-Drought dataset were added with zero counts as demonstrated in the Additional file 1 in **Tab. S1**. Model performance was evaluated, including mean accuracy, F1 score, precision, and recall, which were computed across all taxonomic ranks for both subsets.

## 3 Results

### Alpha and Beta Diversity

Alpha diversity analysis, utilizing the Shannon index as displayed in **Fig. 1 A**, found no significant differences between the ‘Control’ and ‘Drought’ groups. This demonstrates that microbial diversity within individual samples was not significantly impacted by watering regimes. On all taxonomic ranks, no highly abundant taxon was found to be uniquely abundant to drought stress, as only differences between relative abundances between ‘Control’ and ‘Drought’ groups could be observed as shown in Additional file 1 in **Fig. S2**. Beta diversity, assessed via PCoA based on Bray-Curtis dissimilarities as shown in **Fig. 1 B**, yielded insights into the variation between the samples. The watering regime accounted for 6.8 % of the variance and could be clustered into the corresponding irrigation groups. In order to train the ML model to detect drought stress from a variety of soil samples deriving from different isolation sources and crops, the whole dataset was used without subsetting it to specific sample types. For further 16S rRNA-based metagenomic analyses with this dataset and its metadata, *Naylor et al*.*’s* paper itself [31] is referred to, offering interesting insights into the influence of the soil isolation source and the impact of the different crops on the root microbiome.

**Fig. 1.**
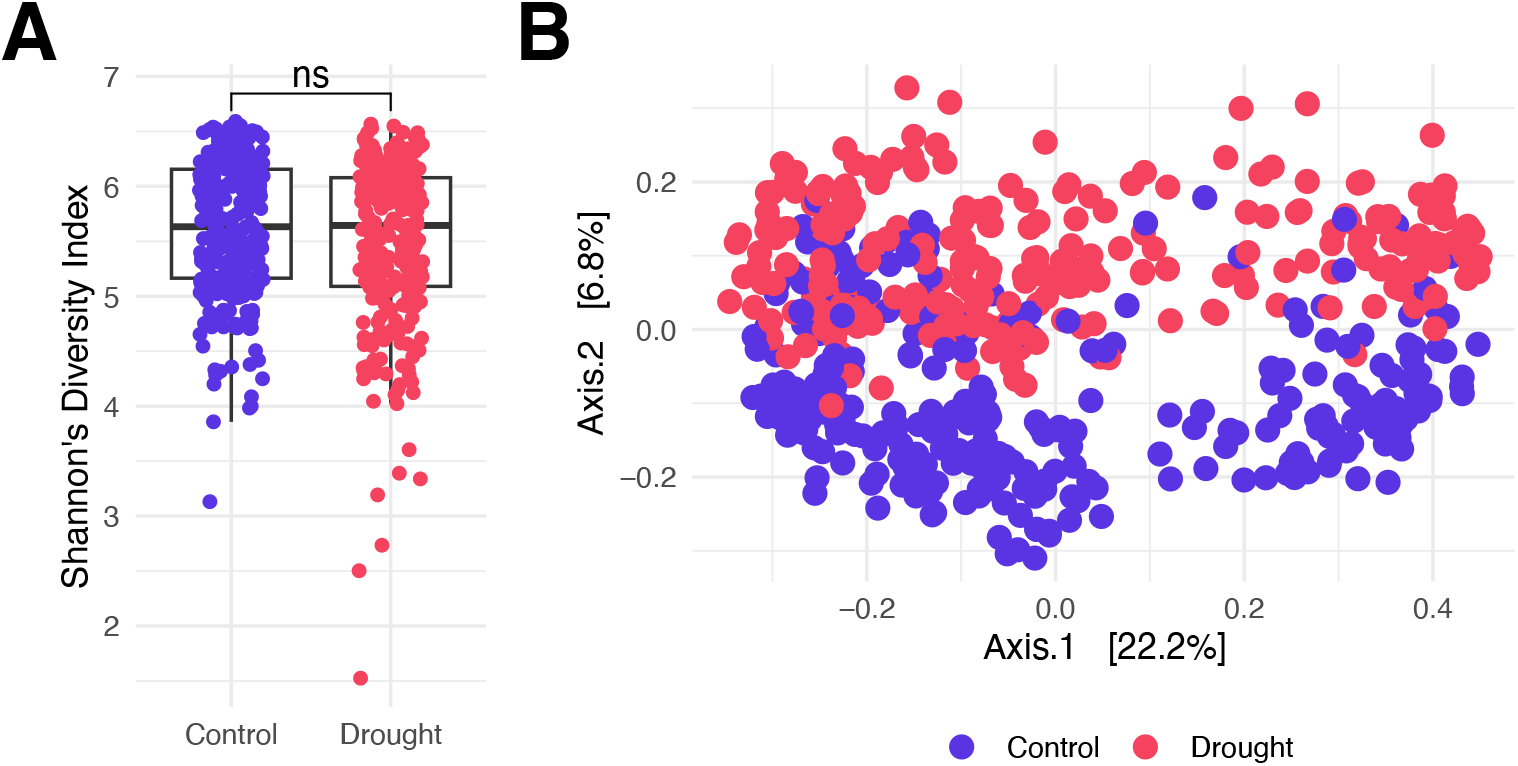
Diversity Plots for the Grass-Drought Dataset. Alpha and beta diversity plots comparing the ‘Control’ (blue) and ‘Drought’ (red) watering regimes. (A) Boxplots of Shannon’s Diversity Index for all samples comparing watering regimes. Significance was determined using a non-parametric Wilcoxon rank sum test (* p *<*0.05, ** p *<*0.01, *** p *<*0.001, **** p *<*0.0001). (B) Principal Coordinate plot using Bray-Curtis dissimilarities colored by the watering regimes.

### Comparative Analysis of DAA Tools

This study’s comprehensive approach to DAA encompassed five distinct methods: DESeq2, ANCOM-BC2, ALDEx2, edgeR, and the non-parametric Wilcoxon rank-sum test (**Fig. 2**). All methods used False Discovery Rate (FDR)-corrected p-values with BH correction and an alpha threshold <0.05. A total of 2,356 ASVs were identified as significantly differentially abundant. Strikingly, 441 ASVs were identified by all five methods, highlighting a core set of differentially abundant taxa. EdgeR and the non-parametric Wilcoxon rank-sum test identified 485 and 318 unique ASVs, respectively. On the other hand, ANCOM-BC2, ALDEx2, and DESeq2 showed consistent results with no or only a small number of uniquely identified ASVs and were therefore used for DAA on all taxonomic ranks. An increased level of consistency between the three tools was visible as displayed in Additional file 1, **Fig. S3**.

**Fig. 2.**
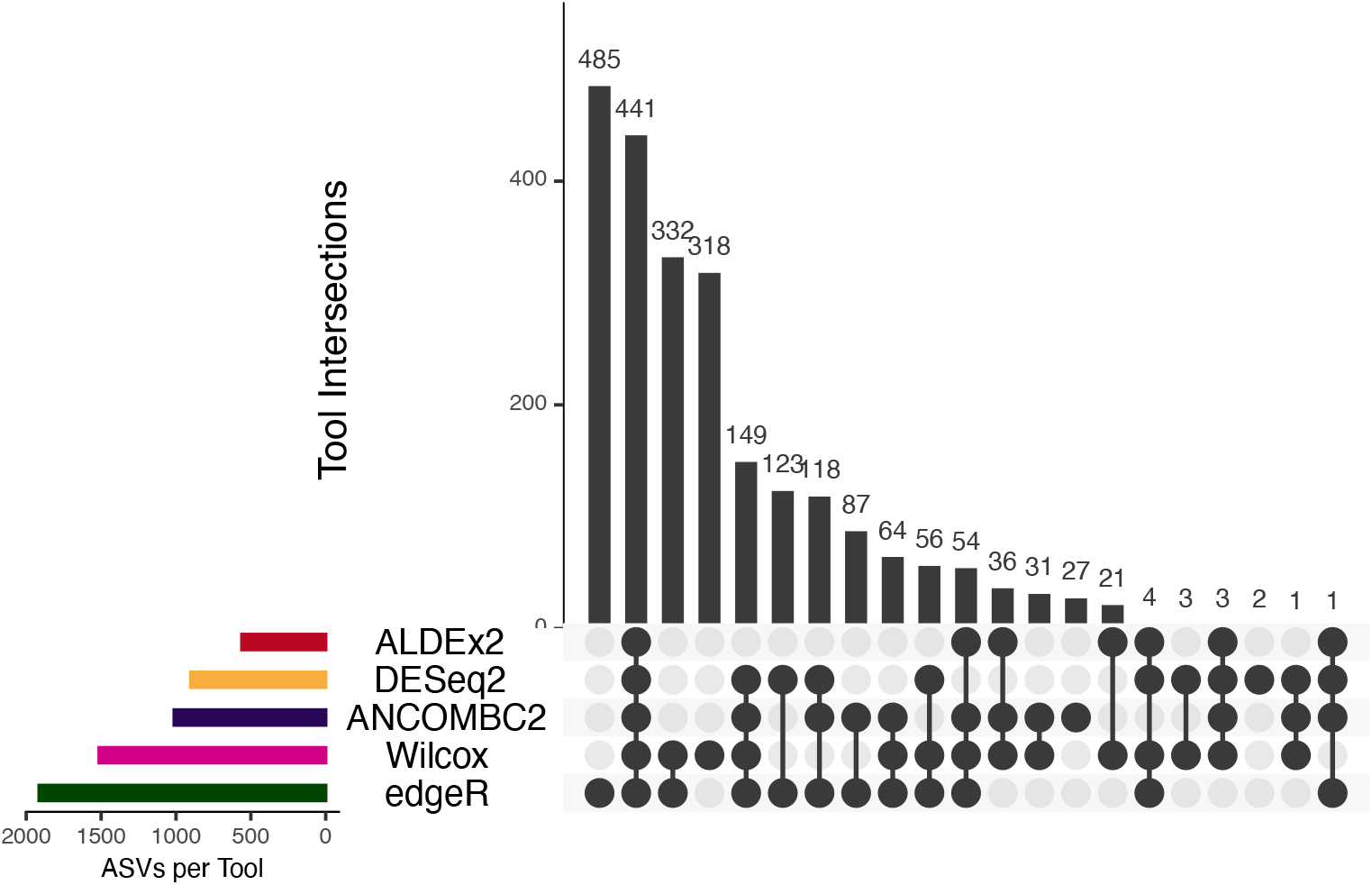
ASV Intersections between different DAA Tools on ASV level of the Grass-Drought Dataset. Upset plots displaying the overlap and uniqueness of significant taxa on ASV level identified by DAA methods (ALDEx2, DESeq2, ANCOM-BC2, non-parametric Wilcoxon ranksum test, and edgeR). The horizontal bars show the total number of taxa for each tool, while the vertical bars show the number of shared taxa between corresponding sets, sorted by the total number of shared taxa. All tools use an alpha threshold of 0.05 for significance.

### The RFC shows remarkable classifying performance across all ranks

Machine learning using the trained RFC demonstrated remarkable performance scores in predicting drought stress in the soil metagenome. **Tab. 1** shows, that across all taxonomic ranks, the RFC consistently delivered exceptional results, with a mean accuracy surpassing 90 %.

**Tab. 1.** Random Forest Classifier Performance of the Grass-Drought Dataset. Table displaying the mean accuracy, F1 score, precision, recall, and AUC of the classifier on different taxonomic ranks of the Grass-Drought dataset, with the best-performing rank for each metric marked in bold.

| Metric | Phylum | Class | Order | Family | Genus |
| --- | --- | --- | --- | --- | --- |
| Accuracy | 0.900 ± 0.025 | 0.911 ± 0.020 | 0.914 ± 0.023 | 0.921 ± 0.017 | **0.923 ± 0.029** |
| F1 score | 0.895 ± 0.029 | 0.906 ± 0.023 | 0.912 ± 0.024 | 0.919 ± 0.017 | **0.921 ± 0.030** |
| Precision | 0.891 ± 0.041 | 0.902 ± 0.032 | 0.890 ± 0.038 | **0.902 ± 0.038** | 0.892 ± 0.041 |
| Recall | 0.899 ± 0.024 | 0.911 ± 0.022 | 0.936 ± 0.022 | 0.939 ± 0.031 | **0.954 ± 0.029** |
| AUC | 0.960 ± 0.020 | 0.960 ± 0.010 | 0.970 ± 0.010 | 0.970 ± 0.010 | **0.980 ± 0.010** |

The genus level proved to be the most effective input, achieving an accuracy of 0.923 ± 0.029, an F1 score of 0.921 ± 0.030, and a recall of 0.954 ± 0.029. Family-level analysis excelled in precision, with a score of 0.902 ± 0.038. Furthermore, the AUC underscored the robust performance of the RFC, with a mean AUC of 0.980 ± 0.010 at the genus level. The corresponding Reciever Operating Characteristic (ROC) curves can be found in Additional file 1 in **Fig. S4**.

### Interpretable ML and DAA as complementary approaches in marker taxa identification

DAA tools (ANCOM-BC2, ALDEx2, DESeq2), and SHAP values emerged as powerful methodologies for investigating taxon enrichment and feature importance, respectively. Both approaches, as displayed in **Fig. 3**, consistently identified taxa responsible for driving differences between the ‘Control’ and ‘Drought’ groups, which can be considered marker taxa for drought stress. Overall, the proportion of matches in the enrichment assignments of DAA tools and SHAP values for all identified taxa ranged from 79.59 % on the order level to 82.65 % on the genus level.

**Fig. 3.**
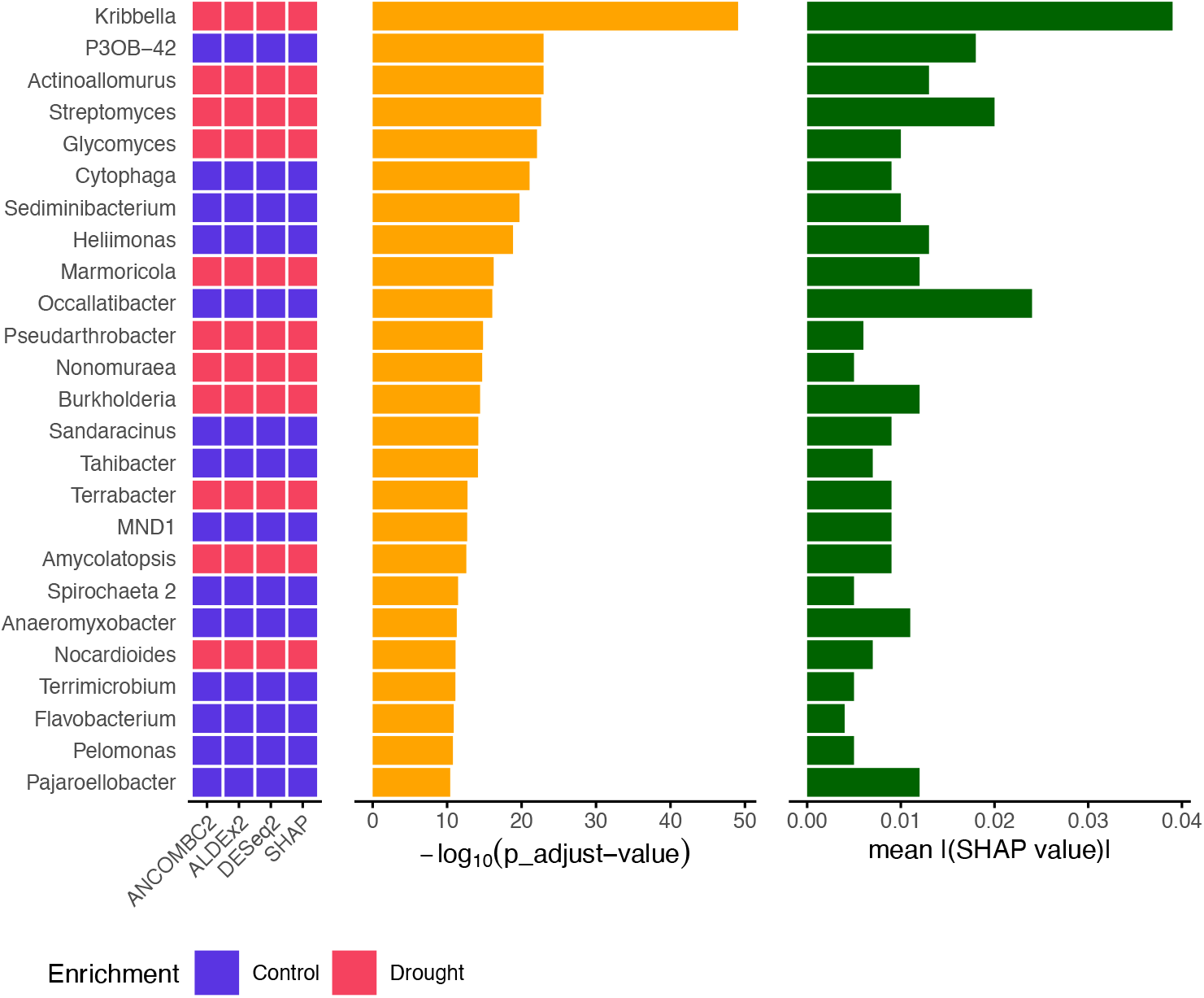
Genera Enrichment, Significance and Importance by DAA Tools and SHAP Values of the Grass-Drought Dataset. Binary heatmap showing the enrichment of the top 25 significant genera from ANCOM-BC2 between ‘Control’ (blue) and ‘Drought’ (red) groups for the three methods used for DAA (DESeq2, ANCOM-BC2, ALDEx2) with an alpha *<*0.05, and SHAP values obtained from the RFC. Corresponding bar plots comparing -log_10_(p adjust) values (orange) and mean(|SHAP value|) (green).

The significance and importance of microbial taxa in the dataset were explored using both DAA significances and SHAP value feature importances of the RFC. Comparing the sorted top 25 genera with adjusted significance levels from ANCOM-BC2 and their corresponding mean absolute SHAP values, it was found that the most significant and also important taxon with more than a two-fold difference to the next taxon was the genus *Kribbella* (**Fig. 3**).

In a direct comparison with the feature importances, it was shown that the order of the taxa identified as important differed greatly from that of ANCOM-BC2 in some genera. This effect was also visible at higher taxonomic ranks (Additional file 1 **Fig. S5** and **Fig. S6**). Especially the genera *Streptomyces* and *Occallatibacter* did not stand out from the results of ANCOM-BC2 but were prioritized considerably higher by their SHAP values.

### The trained RFC generalizes to unseen samples from a different dataset

To assess the generalizability of the trained RFC model, it was applied to a test dataset from *Xu et al*. (2018), focusing on *Sorghum bicolor* root microbiomes subjected to drought stress. First, the trained RFC underwent testing using samples exhibiting advanced drought stress conditions, referred to as the ‘Late Drought’ subset. These samples were expected to demonstrate the most noticeable changes in the relative abundances of the taxa. Stable accuracies across all taxonomic ranks could be detected, as displayed in **Tab. 2**, with a notable improvement towards the family level. The family level achieved the highest accuracy (0.854 ± 0.017), while also excelling in F1 score (0.855 ± 0.021) and precision (0.830 ± 0.029). The order level exhibited the best recall (0.925 ± 0.031).

**Tab. 2.**
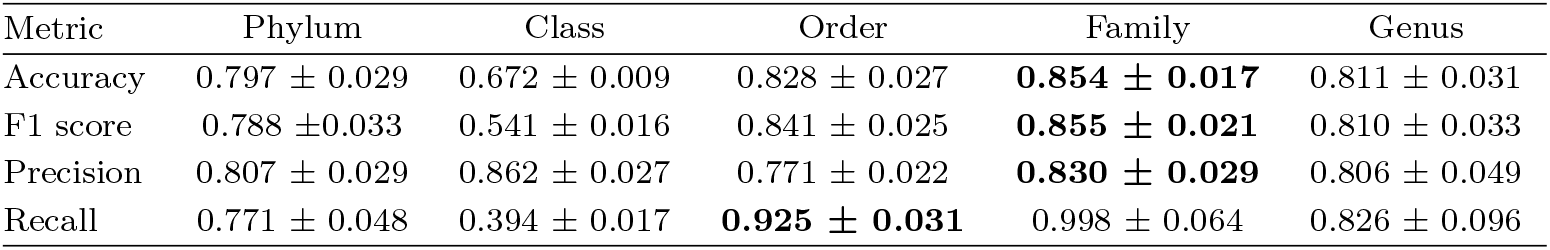
Late Drought Classifier Performance of the Sorghum-Drought Dataset. Table displaying the mean accuracy, F1 score, precision, and recall of the classifier on different taxonomic ranks on the Late Drought subset. The best-performing rank for each metric is marked in bold.

Due to the classifier’s outstanding performance with the ‘Late Drought’ subset, testing extended to another subset containing various drought stress levels, referred to as the ‘Progressive Drought’ subset. This subset contained samples of the complete course of the drought period with associated controls. Here, the order level displayed the best F1 score (0.754 ± 0.024) and recall (0.887 ± 0.018), while the family level yielded the highest accuracy (0.768 ± 0.018) and precision (0.692 ± 0.014) as shown in **Tab. 3**. For both subsets, it was noticeable that the best performance was not observed at the genus level, but at the family or order level.

**Tab. 3.** Progressive Drought Classifier Performance of the Sorghum-Drought Dataset. Table displaying the mean accuracy, F1 score, precision, and recall of the classifier on different taxonomic ranks on the Progressive Drought subset. The best-performing rank for each metric is marked in bold.

| Metric | Phylum | Class | Order | Family | Genus |
| --- | --- | --- | --- | --- | --- |
| Accuracy | $0.678 \pm 0.029$ | $0.613 \pm 0.004$ | $0.750 \pm 0.029$ | **$0.768 \pm 0.018$** | $0.711 \pm 0.044$ |
| F1 score | $0.625 \pm 0.031$ | $0.374 \pm 0.016$ | **$0.754 \pm 0.024$** | $0.753 \pm 0.030$ | $0.697 \pm 0.029$ |
| Precision | $0.628 \pm 0.037$ | $0.614 \pm 0.007$ | $0.656 \pm 0.028$ | **$0.692 \pm 0.014$** | $0.647 \pm 0.055$ |
| Recall | $0.623 \pm 0.038$ | $0.270 \pm 0.015$ | **$0.887 \pm 0.018$** | $0.831 \pm 0.076$ | $0.773 \pm 0.106$ |

## 4 Discussion

This study employed ML to predict the irrigation state of soil samples based on their microbial community composition and aimed to discover specific marker taxa for drought stress. In terms of alpha and beta diversity analysis, it is crucial to note that drought stress primarily influenced the relative abundance of taxa rather than causing a complete abolishment or appearance of certain taxa [52], which aligns with the study’s focus on the ‘Control’ and ‘Drought’ labels. This suggests that the classifier was trained by emphasizing variations in taxon abundance rather than focusing on the presence or absence of specific taxa.

Regarding the comparison of interpretable ML with conventional DAA tools for marker taxa identification, the focus was on finding the most suitable DAA tools for the Grass-Drought dataset from a variety of popular methods, namely ALDEx2, ANCOM-BC2, DESeq2, edgeR, and the non-parametric Wilcoxon rank-sum test. This approach is generally recommended for DAA, as it is not possible to find the true number of significant taxa in real-world data sets like it is the case with mock data [53, 54]. The used DAA methods made different assumptions about the data distribution [53]. For instance, DESeq2 and edgeR assume a negative binomial distribution, while ALDEx2 and ANCOM-BC2 assume a Gaussian distribution. The non-parametric Wilcoxon rank-sum test, on the other hand, does not make any distribution assumptions.

On the ASV level, out of 3,276 total assigned ASVs, 71.9 % were identified as significant by at least one of the five DAA methods. However, only 13.46 % of these significant ASVs were detected by all five methods, suggesting a substantial proportion of ASVs being uniquely identified by specific tools, possibly indicating false discoveries [53]. Specifically, edgeR and the Wilcoxon rank-sum test uniquely identified a high number of ASVs not detected by other tools, which can be an indicator for many false discoveries and unreliable results [55].

In contrast, ANCOM-BC2, DESeq2, and ALDEx2 exhibited more reliable results, with fewer uniquely identified ASVs. These findings are in line with previous studies that have highlighted the reliability of these three methods in controlling the FDR [53, 54, 56, 57]. Out of these three tools, ANCOM-BC2 was selected as the DAA method to compare its results directly with those of interpretable ML, as it showed the most overlap in detected ASVs with the other two selected DAA tools.

For interpretable ML, a RFC was chosen as it is recognized as a top-performing classifier for handling high-dimensional and sparse data, such as metagenomic datasets with hundreds to thousands of features and non-linear relationships between features and the target variable [58–62].

The classifier, trained on a dataset containing soil samples from a variety of soil isolation sources, crops, and drought stress levels, exhibited exceptional performance across all taxonomic ranks, directly addressing objective a) of this study. As the taxonomic rank descended from higher (e.g., phylum) to lower levels (e.g., genus), the granularity and resolution of the features increased. At the genus level, which represented the lowest taxonomic rank, the classifier exhibited the highest mean accuracy of 92.3 %, as well as the highest F1 score, recall, and AUC, demonstrating its effectiveness in capturing true positive instances. While there was a slight increase in overall accuracy from the phylum to the genus level, the genus level provided more specific insights into microbial diversity and potential marker taxa for drought stress.

In the exploration of marker taxa for drought stress in the soil metagenome, interpretable ML was employed using SHAP values in the nested CV of each taxonomic rank. Enrichment and feature importance results were compared with the output of the three most suitable DAA tools for this dataset. At higher taxonomic ranks, the agreement between DAA enrichment and SHAP value contribution was less definitive. Among the DAA tools, taxa were often not classified as significant by all three tools, as seen with Bacteroidota at the phylum level. Similarly, SHAP values did not always provide clear results between the loops of the nested CV, making it challenging to assign enrichment to either ‘Control’ or ‘Drought’, as observed with Firmicutes at the phylum level. In some cases, the results from DAA and interpretable ML differed, such as with Verrucomicrobiae and Armanimonadota at the phylum level, which were classified as enriched in ‘Control’ by DAA tools and enriched under ‘Drought’ by SHAP. According to the literature, both phyla were found to be more enriched under irrigation, but the same study concluded that both taxa have the potential to assist plants under drought conditions [63]. However, at lower, more specific ranks such as family and genus levels, all enrichment information among the top 25 taxa is consistent. This consistency highlights that SHAP values can be equally useful for the discovery of specific marker taxa under stress conditions, effectively fulfilling objective b).

Furthermore, the rankings of taxa between DAA and ML approaches were compared. While the order of significant taxa differed, the genus *Kribbella* consistently emerged as most significant and important, displaying a two-fold increase compared to the next relevant genus. Although being a poorly studied genus, *Kribbella* has shown potential in promoting plant growth [64, 65], making it a promising marker taxon for drought stress.

Additionally, in the direct comparison of significances and feature importances, certain taxa were detected with greater prominence, as evidenced by a peak in their mean absolute SHAP value compared to the significance assigned by the DAA analysis, like the genera *Streptomyces* and *Occallatibacter. Streptomyces*, a dominant genus in soil microbiomes, has been associated with drought stress and plant health in dry environments [66–68]. The genus *Occallatibacter* showed depletion under drought conditions and was considered an important feature for the prediction of drought stress, although further research is needed to understand its specific impact on soil metagenomes under drought, as it has only been observed under heat stress and no-stress conditions [69, 70].

SHAP values and DAA tools use different underlying approaches for the identification of important or significant taxa. In the context of this study, it is not possible to determine which approach is more suitable, but the overall results suggest that both methods provide important information for the identification of marker taxa. Therefore, these approaches should be seen as complementary rather than interchangeable, with each providing valuable insights into metagenomic data analysis.

To evaluate the generalization capabilities of this study’s classifier, its performance was tested on another drought stress dataset. The classifier’s performance was assessed with samples undergoing several weeks of drought stress as the most impactful differences were expected between the two watering regimes. The Late Drought subset exhibited an accuracy score of 0.854 ± 0.017 at the family level. Therefore, the classifier’s effectiveness and robustness across the entire spectrum of drought stress levels of the Sorghum-Drought dataset was explored by predicting drought stress in the Progressive Drought subset. Remarkably, the results consistently demonstrated the classifier’s outstanding performance in both scenarios. The Progressive Drought subset achieved an accuracy score of 0.768 ± 0.018 at the family level, indicating the model’s reliability in classifying drought stress regardless of the drought stress level involved.

In contrast to the Grass-Drought dataset, where the best performance was achieved at the genus level as the lowest taxonomic rank with the highest granularity, the subsets of the Sorghum-Drought test dataset did not yield the best classification results on this rank. The best performance was observed on the order and family levels. This can be attributed to the inherent sparsity on the genus level due to the addition of taxa with zero counts to create feature tables for the prediction with equal feature inputs, as displayed in the Additional file 1 in **Tab. S1**.

These results emphasize the classifier’s adaptability across diverse drought stress conditions, reinforcing its utility as a valuable tool for drought stress classification, in line with the objectives outlined in part c) of this study. Even though the classifier was trained with a dataset containing 16S rRNA metagenomic data of different drought stress levels, soil isolation sources, and a variety of plants, the approach might vary based on input data from other sequencing regions or plants that the classifier was not trained on. Such differences may emerge due to differences in the estimation of microbial diversity [71, 72]. The effect of the soil isolation source on the prediction also offers scope for further investigations to improve the classifier’s predictive capabilities and potentially develop a classifier tailored to specific soil isolation sources [73]. Expanding the sample size could further enhance the classifier’s generalizability, as a more extensive representation of taxa would increase the likelihood of encountering taxa abundances present in unknown samples during new predictions. However, when dealing with large datasets, selecting a comprehensive range of core taxa is recommended. By selecting the most important taxa as features [74] and training the classifier accordingly, the introduction of sparsity in the feature tables when predicting new data can be prevented. Furthermore, including samples from different locations introduces another dimension, as variations in microbial composition across diverse geographic locations and climates [75] can impact the classifier’s performance. With the upcoming possibilities of 16S rRNA long-read sequencing, it is recommended to create a location- and long-read sequencing-based classifier to generate an individual classifier with a reduced bias. Further evaluation with different ML models, more data, and subsequent feature selection seem interesting applications for future research.

## 5 Conclusions

In conclusion, with the ongoing threat of extreme weather events, notably droughts [76, 77], it is indispensable to explore innovative methods for understanding the impact of the soil microbiome on agriculture and ecosystems. The primary accomplishment of this study is the creation of a location-based classifier for drought stress in the soil metagenome. Demonstrating remarkable generalization capabilities, the classifier assesses drought stress across various drought stress levels and is applicable to different grasses.

The application of this study’s generalized ML model extends beyond the classification of drought stress, facilitating precision agriculture, including the optimization of irrigation strategies. Further, in microbe-assisted plant breeding programs, the discovery of marker taxa for drought stress using interpretable ML with SHAP values provides farmers and breeders with valuable insights for the definition of microbial strains for targeted bioinoculation approaches. Here, a deeper understanding of the plant growth-promoting functions associated with drought stress-related taxa holds promise for future advancements. This knowledge could play a pivotal role in enhancing plant adaptation to drought stress, strengthening the plant immune system against yield losses, and reducing susceptibility to pathogens [78].

## 6 List of abbreviations

ASVs: Amplicon Sequence Variants
AUC: Area Under the Curve
BH: Benjamini-Hochberg
CV: Cross-Validation
DAA: Differential Abundance Analysis
FDR: False Discovery Rate
ML: Machine Learning
PCoA: Principal Coordinate Analysis
RDP: Ribosomal Database Project
RFC: Random Forest Classifier
ROC: Reciever Operating Characteristic
SHAP: SHapley Additive ExPlanation
SynComs: Synthetic Communities

## Supplementary Information

**Additional file 1**

- File name: Additional file 1
- File format: .pdf
- Title of data: Supplementary Figures and Tables
- Description of data: **Fig. S1:** Weekly Watering Scheme of the Sorghum-Drought Test Dataset. Watering scheme for the three groups ‘Control’ (C), ‘Pre-Flowering Drought’ (PrD), and ‘Post-Flowering Drought’ (PoD). Dashed lines indicate watering and white cells indicate no watering. The test dataset was subset into the ‘Progressive Drought’ subset (red) and ‘Late Drought’ subset (blue). **Tab. S1:** Feature Table Pruning between the Datasets. Table displaying differences in the total number of different taxa between the Grass-Drought dataset as the dataset the RFC has been trained on and the Sorghum-Drought dataset as the test dataset. A high number of taxa added with zero counts in order to bring the feature tables of the test dataset into a suitable format for prediction also increases the level of potential sparsity. **Fig. S2:** Relative Abundances per Rank of the Grass-Drought Dataset. Bar plots displaying the relative abundance of the top 10 taxa between the ‘Control’ and ‘Drought’ groups on (A) Phylum, (B) Class, (C) Order, (D) Family, and (E) Genus level in alphabetical order. **Fig. S3:** Significant Taxa Intersections between DAA Tools per Rank of the Grass-Drought Dataset. Upset plots displaying the overlap and uniqueness of significant taxa identified by the three DAA methods ‘ALDEx2’, ‘DESeq2’, and ‘ANCOM-BC2’ on (A) Phylum, (B) Class, (C) Order, (D) Family, and (E) Genus level. The horizontal bars show the total number of taxa for each tool, while the vertical bars show the number of shared taxa between corresponding sets, sorted by the total number of shared taxa. All tools use an alpha threshold of 0.05 for significance. **Fig. S4:** ROC Curves per Rank of the Grass-Drought Dataset. Receiver Operating Characteristic (ROC) curve at (A) Phylum, (B) Class, (C) Order, (D) Family, and (E) Genus level, showing the area under the curve (AUC) for each fold of the nested cross-validation. The ROC curve displays the best model for each fold and the mean AUC. **Fig. S5:** Taxon Enrichment, Significance, and Importance by DAA Tools and SHAP Values of the Grass-Drought Dataset. Binary heatmap showing the enrichment of the top significant genera from ANCOM-BC2 on (A) Phylum and (B) Class level between ‘Control’ (blue) and ‘Drought’ (red) groups for the three methods used for DAA (DESeq2, ANCOM-BC2, ALDEx2) with an alpha <0.05, and SHAP values obtained from the RFC. Empty cells display no significant enrichment. Corresponding bar plots comparing -log_10_(p adjust) values (orange) and mean(|SHAP value|) (green). **Fig. S6:** Taxon Enrichment, Significance, and Importance by DAA Tools and SHAP Values of the Grass-Drought Dataset. Binary heatmap showing the enrichment of the top significant genera from ANCOM-BC2 on (A) Order and (B) Family level between ‘Control’ (blue) and ‘Drought’ (red) groups for the three methods used for DAA (DESeq2, ANCOM-BC2, ALDEx2) with an alpha <0.05, and SHAP values obtained from the RFC. Empty cells display no significant enrichment. Corresponding bar plots comparing -log_10_(p adjust) values (orange) and mean(|SHAP value|) (green).

## Declarations

### 6.1 Ethics approval and consent to participate

The study incorporated 16S rRNA metagenomic data from plants and soil that is publicly available and was conducted in accordance with the present ethical guidelines as of the date of publication.

### 6.2 Consent for publication

Not applicable

### 6.3 Availability of data and material

The datasets supporting the conclusions of this article are available in the NCBI Short Read Archive under the BioProjectID PRJNA369551 (‘Grass-Drought’ dataset), and PRJNA435634 (‘Sorghum-Drought’ dataset). All analysis scripts are available on GitHub (https://github.com/Computomics/SoilMicrobiomeDroughtML).

### 6.6 Authors’ contributions

MH conceived, designed, and performed the analysis with inputs from SP, RD, CW, SJS, and JB. MH drafted the manuscript and it was reviewed and edited by all authors. All authors read and approved the final manuscript.

## 6.7 Acknowledgements

Not applicable

## 6.8 Authors’ information (optional)

Not applicable

