## Additional file 1 for "Interpretable Machine Learning Decodes Soil Microbiome’s Response to Drought Stress"

Supplementary Figures and Tables

Michelle Hagen[1], Rupashree Dass[1], Cathy Westhues[1], Jochen Blom[2], Sebastian J Schultheiss[1], Sascha Patz[1]

---

[1] Computomics GmbH
Eisenbahnstraße 1
Tübingen, 72072
Baden-Württemberg, Germany

[2] Bioinformatics & Systems Biology
Justus Liebig University Gießen
Heinrich-Buff-Ring 58
Gießen, 35390
Hesse, Germany

This document includes the following supporting information:

- **Fig. S1:** Weekly Watering Scheme of the Sorghum-Drought Test Dataset
- **Tab. S1:** Feature Table Pruning between the Datasets
- **Fig. S2:** Relative Abundances per Rank of the Grass-Drought Dataset
- **Fig. S3:** Significant Taxa Intersections between DAA Tools per Rank of the Grass-Drought Dataset
- **Fig. S4:** ROC Curves per Rank of the Grass-Drought Dataset
- **Fig. S5 + S6:** Taxon Enrichment, Significance and Importance by DAA Tools and SHAP Values of the Grass-Drought Dataset

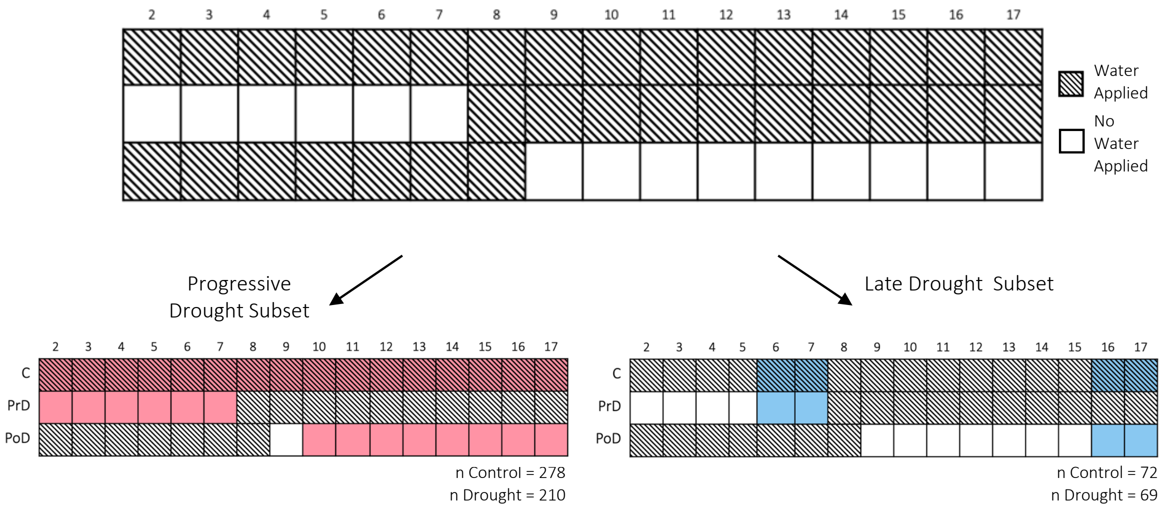


**Fig. S1: Weekly Watering Scheme of the Sorghum-Drought Test Dataset.** Watering scheme for the three groups 'Control' (C), 'Pre-Flowering Drought' (PrD), and 'Post-Flowering Drought' (PoD). Dashed lines indicate watering and white cells indicate no watering. The test dataset was subset into the 'Progressive Drought' subset (red) and 'Late Drought' subset (blue).

**Tab. S1: Feature Table Pruning between the Datasets.** Table displaying differences in the total number of different taxa between the Grass-Drought dataset as the dataset the RFC has been trained on and the Sorghum-Drought dataset as the test dataset. A high number of taxa added with zero counts in order to bring the feature tables of the test dataset into a suitable format for prediction also increases the level of potential sparsity.

| Rank | Phylum | Class | Order | Family | Genus |
|---|---|---|---|---|---|
| Number of taxa grass-drought | 26 | 60 | 131 | 186 | 330 |
| Number of taxa sorghum-drought | 33 | 69 | 148 | 186 | 311 |
| Matching between both | 25 | 54 | 111 | 148 | 198 |
| Added with zero counts | 1 | 6 | 20 | 38 | 132 |

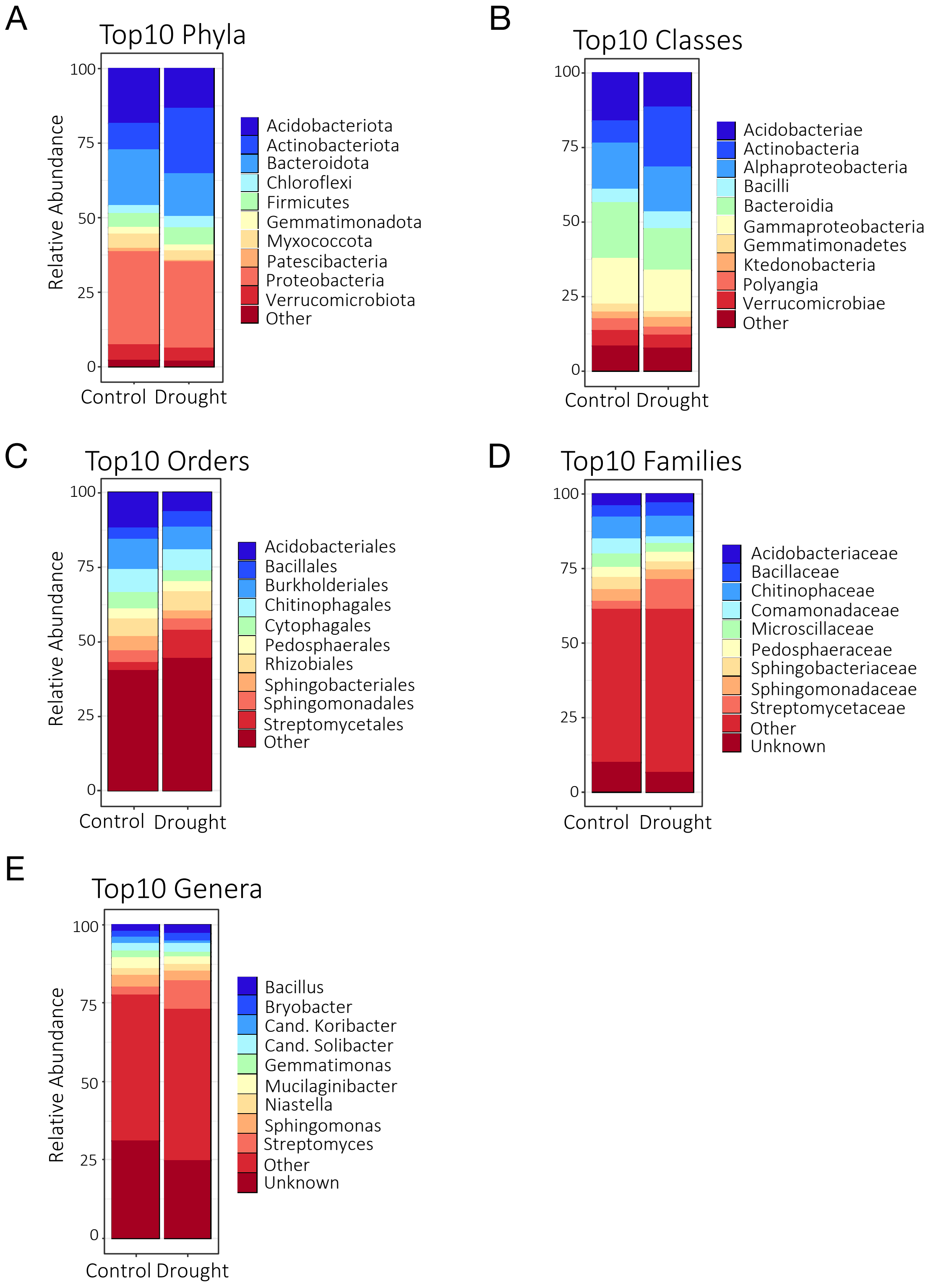


**Fig. S2: Relative Abundances per Rank of the Grass-Drought Dataset.** Bar plots displaying the relative abundance of the top 10 taxa between the 'Control' and 'Drought' groups on (A) Phylum, (B) Class, (C) Order, (D) Family, and (E) Genus level in alphabetical order.

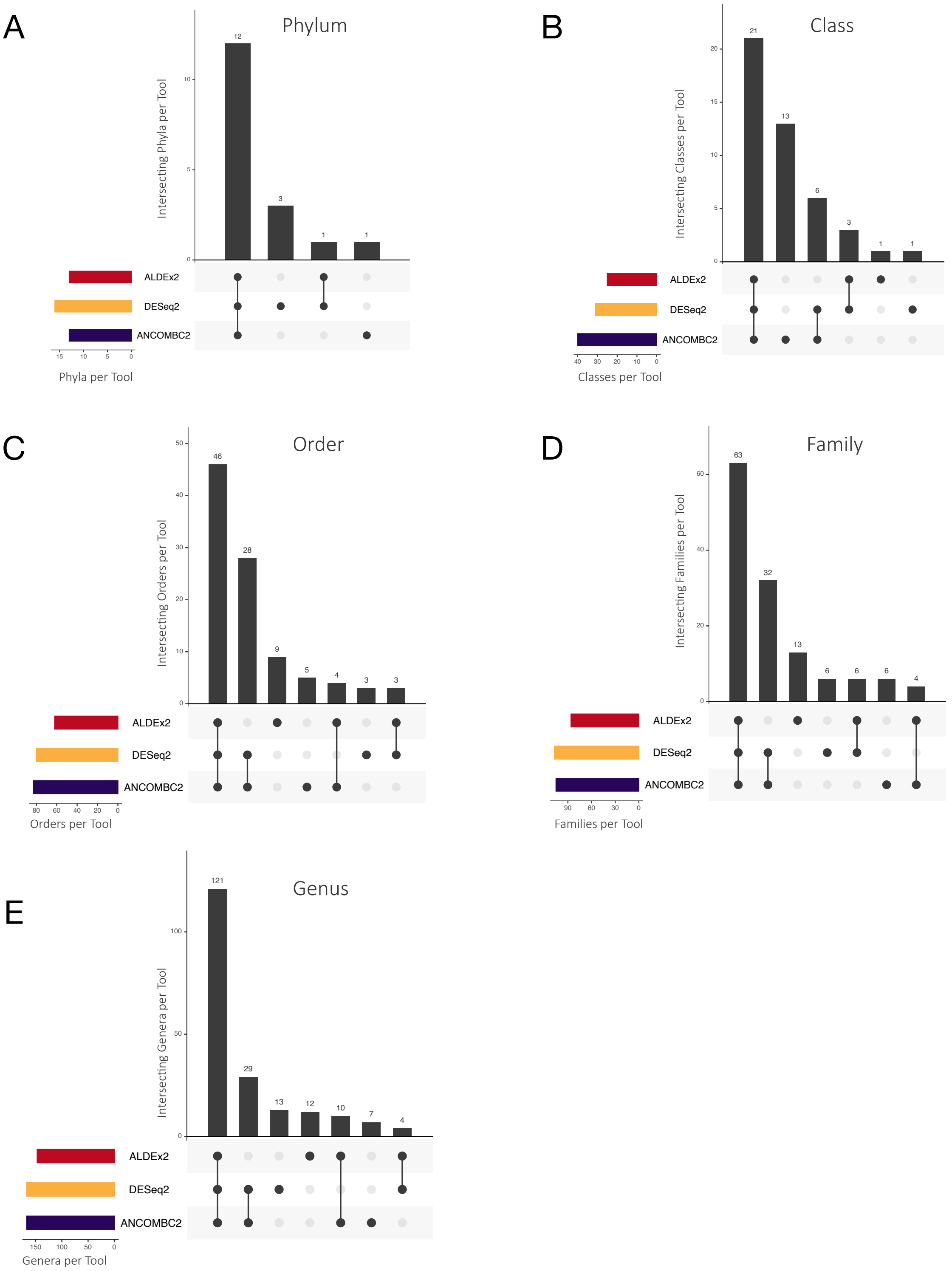


**Fig. S3: Significant Taxa Intersections between DAA Tools per Rank of the Grass-Drought Dataset.** Upset plots displaying the overlap and uniqueness of significant taxa identified by the three DAA methods 'ALDEx2', 'DESeq2', and 'ANCOM-BC2' on (A) Phylum, (B) Class, (C) Order, (D) Family, and (E) Genus level. The horizontal bars show the total number of taxa for each tool, while the vertical bars show the number of shared taxa between corresponding sets, sorted by the total number of shared taxa. All tools use an alpha threshold of 0.05 for significance.

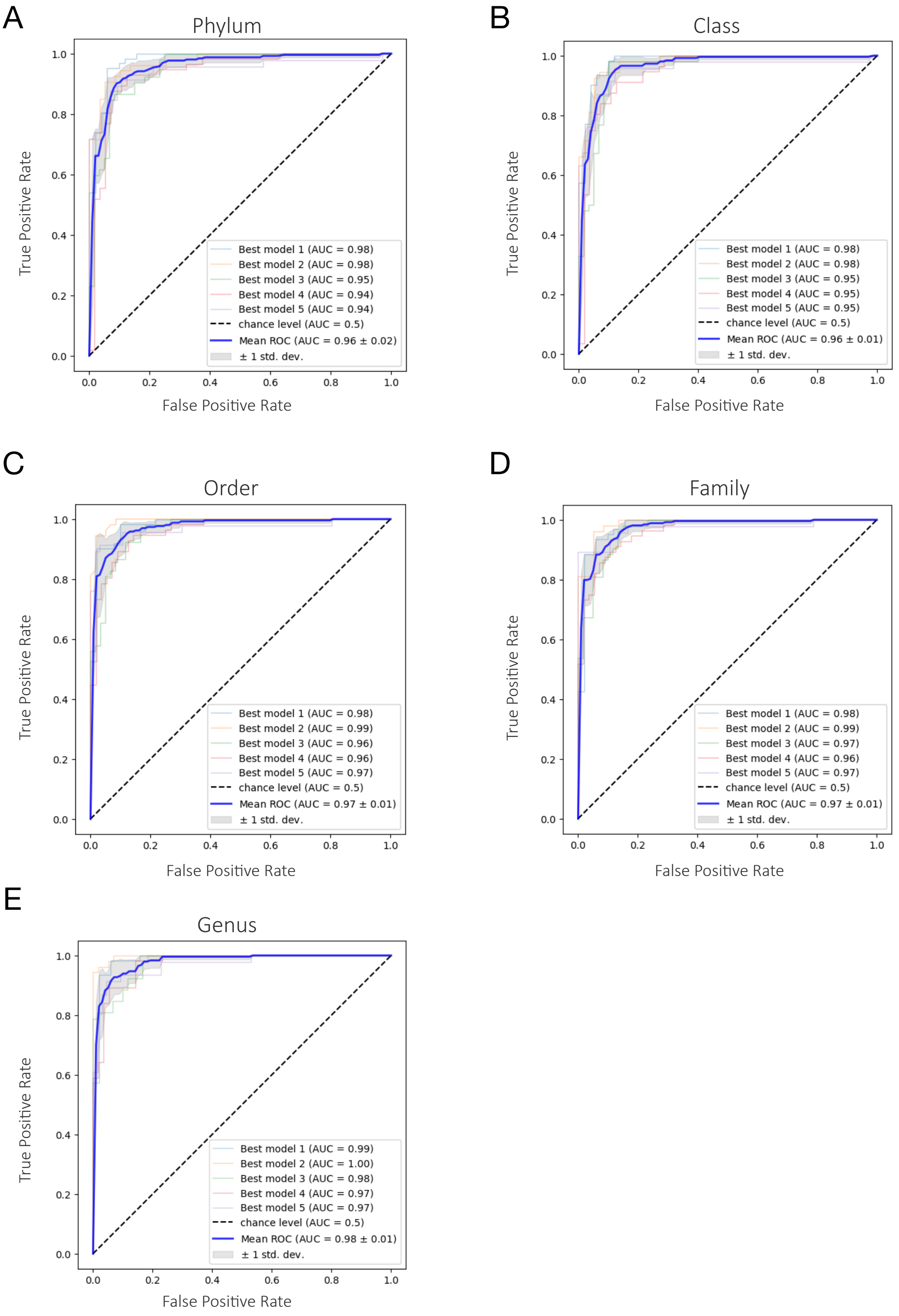


**Fig. S4: ROC Curves per Rank of the Grass-Drought Dataset.** Receiver Operating Characteristic (ROC) curve at (A) Phylum, (B) Class, (C) Order, (D) Family, and (E) Genus level, showing the area under the curve (AUC) for each fold of the nested cross-validation. The ROC curve displays the best model for each fold and the mean AUC.

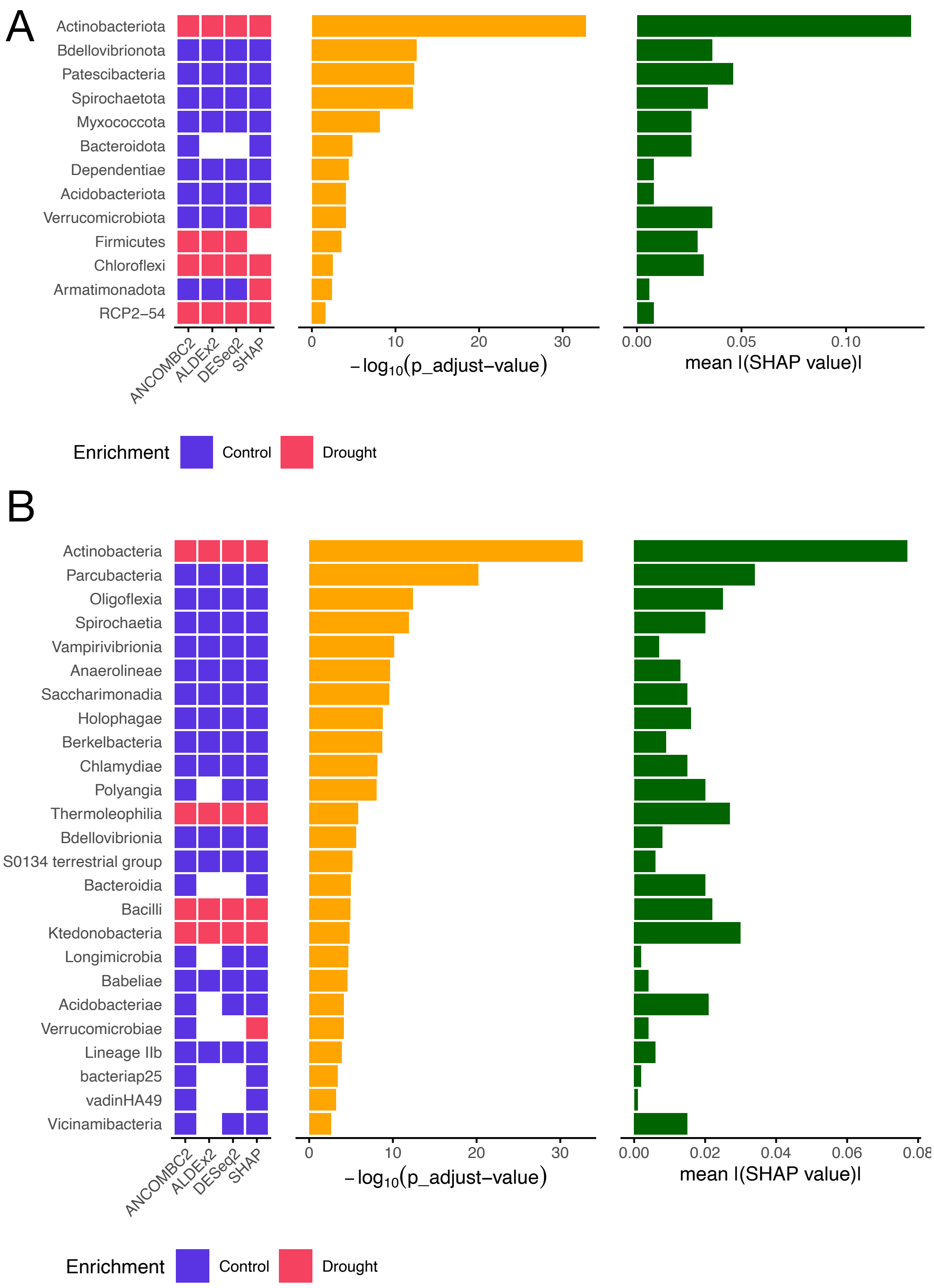


**Fig. S5: Taxon Enrichment, Significance and Importance by DAA Tools and SHAP Values of the Grass-Drought Dataset.** Binary heatmap showing the enrichment of the top significant taxa from ANCOM-BC2 on (A) Phylum and (B) Class level between 'Control' (blue) and 'Drought' (red) groups for the three methods used for DAA (DESeq2, ANCOM-BC2, ALDEx2) with an alpha $<0.05$, and SHAP values obtained from the RFC. Empty cells display no significant enrichment. Corresponding bar plots comparing -$\log_{10}$(p_adjust) values (orange) and mean(|SHAP value|) (green).

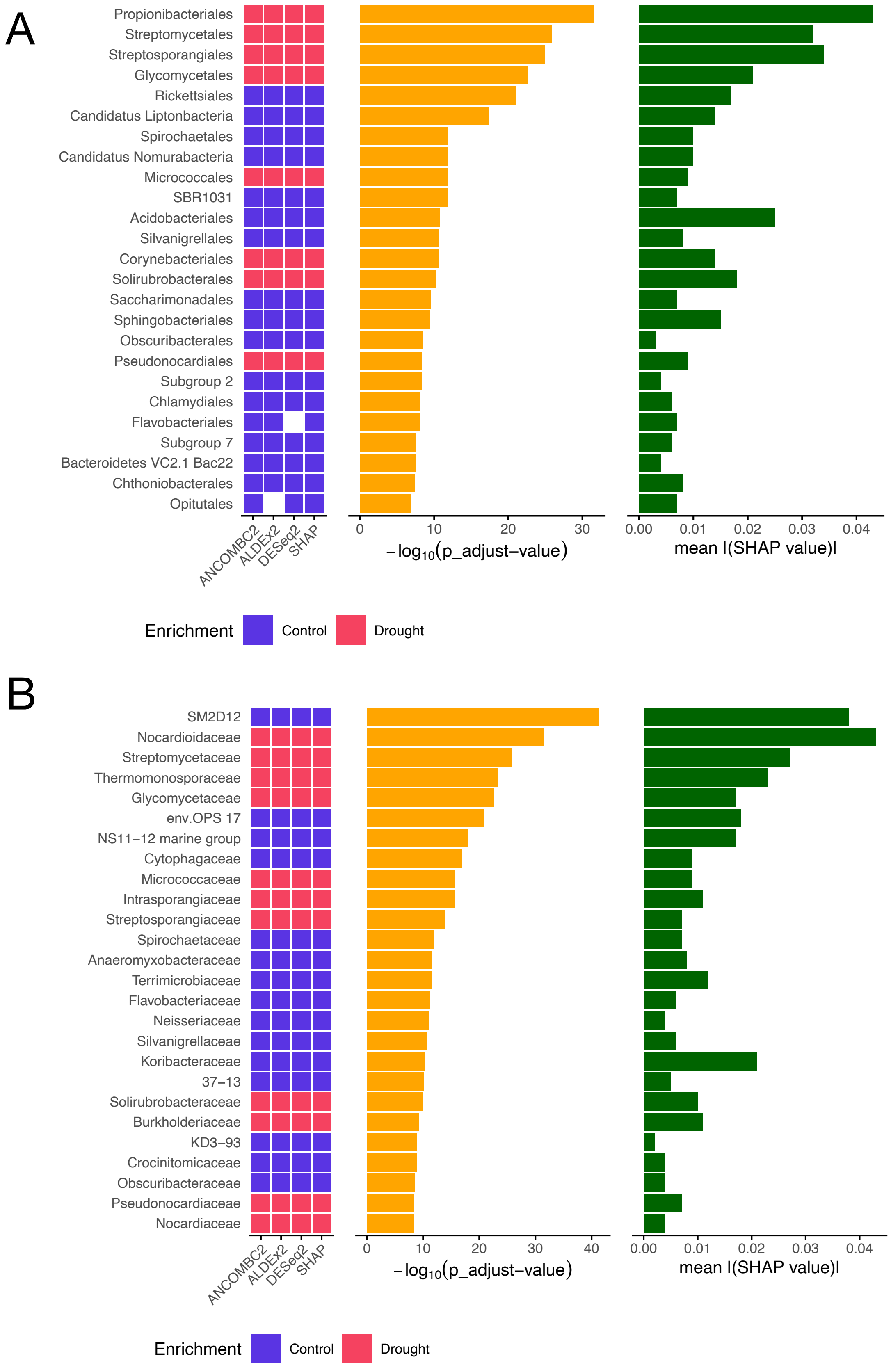


**Fig. S6: Taxon Enrichment, Significance and Importance by DAA Tools and SHAP Values of the Grass-Drought Dataset.** Binary heatmap showing the enrichment of the top significant taxa from ANCOM-BC2 on (A) Order and (B) Family level between 'Control' (blue) and 'Drought' (red) groups for the three methods used for DAA (DESeq2, ANCOM-BC2, ALDEx2) with an alpha $<0.05$, and SHAP values obtained from the RFC. Empty cells display no significant enrichment. Corresponding bar plots comparing -$\log_{10}$(p_adjust) values (orange) and mean(|SHAP value|) (green).
